# Dark period transcriptomic and metabolic profiling of two diverse *Eutrema salsugineum* accessions

**DOI:** 10.1101/163980

**Authors:** Jie Yin, Michael J. Gosney, Brian P. Dilkes, Michael V. Mickelbart

## Abstract

*Eutrema salsugineum* is a model species for the study of plant adaptation to abiotic stresses. Two accessions of *E. salsugineum*, Shandong (SH) and Yukon (YK), exhibit contrasting morphology, biotic, and abiotic stress tolerance. Transcriptome and metabolic profiling from tissue samples collected during the dark period were used to investigate the molecular and metabolic bases of these contrasting phenotypes. RNA sequencing identified 17,888 expressed genes, of which 157 were not in the published reference genome and 65 were detected for the first time. Differential expression was detected for only 31 genes. The RNA sequencing data contained 14,808 single nucleotide polymorphisms (SNPs) in transcripts, 3,925 of which are newly identified. Among the differentially expressed genes, there were no obvious candidates for the physiological or morphological differences between SH and YK. Metabolic profiling indicated that YK accumulates free fatty acids and long-chain fatty acid derivatives as compared to SH; whereas sugars are more abundant in SH. Metabolite levels suggest that carbohydrate and respiratory metabolism, including starch degradation, is more active during the first half of the dark period in SH. These metabolic differences may explain the greater biomass accumulation in YK over SH. The accumulation of 56% of the identified metabolites was lower in F_1_ hybrids than the mid-parent averages and the accumulation of 17% of the metabolites in F_1_ plants transgressed the level in both parents. Concentrations of several metabolites in F_1_ hybrids agree with previous studies and suggest a role for primary metabolism in heterosis. The improved annotation of the *E. salsugineum* genome and newly-identified high-quality SNPs will permit accelerated studies using the standing variation in this species to elucidate the mechanisms of its diverse adaptations to the environment.

## Introduction

*Eutrema salsugineum* (formerly *Thellungiella halophila*) is a model species for the study of plant stress tolerance (Amtmann, 2009; Bressan et al., 2001; Griffith et al., 2007; Oh et al., 2010; Orsini et al., 2010; Pilarska et al., 2016; Taji et al., 2004; Volkov et al., 2003; Wong et al., 2005). The two most commonly studied accessions, Shandong (SH) and Yukon (YK), are native to the Yellow River region of China (Bressan et al., 2001; Inan et al., 2004) and the Yukon territories of Canada (Wong et al., 2005), respectively. These accessions contrast in cold tolerance (Lee et al., 2012), water stress tolerance (MacLeod et al., 2014; Xu et al., 2014), and disease resistance (Yeo et al., 2014). In response to water stress for example, YK accumulates more cuticular wax (Xu et al., 2014), exhibits delayed wilting due to higher leaf water content, and maintains a higher leaf area, as compared to SH (MacLeod et al., 2014). Some differences in adaptive mechanisms have been linked to metabolism, such as a more pronounced increase in fructose and proline content after cold acclimation in YK compared to SH (Lee et al., 2012).

RNA-seq has been used to identify differentially expressed genes (DEGs) that contribute to genotypic variation in response to physiological conditions (Bazakos et al., 2012; Huang et al., 2012a; Huber-Keener et al., 2012; Niederhuth et al., 2013; Stein and Waters, 2012; Wang et al., 2009). When comparing expression between polymorphic accessions of a species, RNA-seq allows for the simultaneous identification and quantification of single nucleotide polymorphisms (SNPs) (Nielsen et al., 2011; Wang et al., 2009). These data can later be utilized as genetic markers for linkage mapping (Chen et al., 2011; Hancock et al., 2011; Hyten et al., 2009; Lisec et al., 2008; Trick et al., 2012) and to assess allele-specific expression (Oshlack et al., 2010; Pickrell et al., 2010).

Plant metabolic profiling permits the simultaneous measurement of multiple intermediates and the end-products of biochemical pathways. Similar to RNA-seq, metabolic profiling can be used to investigate the metabolic and physiological status of biological systems (Fiehn et al., 2000) and may provide biochemical bases for differences in growth and physiology (Meyer et al., 2007). Metabolite profiling has revealed correlations between particular metabolites and growth in *Arabidopsis thaliana* (Meyer et al., 2007), but it is not clear whether these metabolites are more universally linked to heterosis.

The use of combined transcriptome and metabolome analyses have been successfully utilized to identify novel genes and enzymes in plant metabolite biosynthesis pathways (Boke et al., 2015; Sumner et al., 2015). In addition, this approach has also provided insights into pathways that are affected by gene mutation (Etalo et al., 2013; Masclaux-Daubresse et al., 2014; Page et al., 2016; Satou et al., 2014), that are related to responses to biotic stresses (Gurkok et al., 2015; Liu et al., 2016), and abiotic stresses (Bielecka et al., 2014; Hamanishi et al., 2015).

Together, transcriptome and metabolite profiling provide complementary experimental evidence to guide the construction of rational hypotheses for the biochemical basis of variation in growth. The goal of this study is to identify the metabolic and transcription bases for the growth differences between the two *E. salsugineum* accessions. We utilized contrasting genotypes to identify genetic differences, expression divergence and metabolic compounds associated with observed phenotypic variation. We obtained gene expression and metabolite concentration data from SH and YK accessions during the dark period by RNA-seq and gas chromatography mass spectrometry (GC/MS), respectively. We utilized RT-PCR to identify DEGs implicated by our RNA-seq experiment, validating 23 of 25 candidates. By mining the RNA-seq experiment, we validated previously identified SNPs and discovered additional SNPs that differentiate these *E. salsugineum* accessions. We propose that observed differences in metabolite accumulation could contribute to differences in biomass.

## Materials and Methods

### Plant material and growth conditions

Seeds of *E. salsugineum* SH and YK accessions were obtained from individual selfed plants. To generate SH x YK F_1_ seed, closed flower buds of YK plants were manually opened, anthers removed, and pollen from SH plants was applied to the stigma. Multiple crosses (ca. 20) were made on a single plant, which was kept in isolation until seed set. Seeds from this plant were used for F_1_ experiments.

Seeds were stratified in the dark for 10 days at 4°C then sown in a 4:1 mix of Fafard 52 Mix (transcriptome and first metabolome experiments) or Fafard 2 Mix (second metabolome experiment) soilless media (Conrad Fafard, Inc., Agawam, MA, USA) and Turface calcined clay (Profile Products LLC., Buffalo Grove, IL, USA). Plants were grown in 50 mL conical tubes (USA Scientific, Inc., Ocala, FL, USA). A 0.5 cm hole was drilled at the bottom of each tube where seeds were sown. Tubes were then closed with a mesh cap and placed cap-side-down to allow for sub-irrigation. Tubes were placed in a mist room for 12 days, at which point seedlings had reached the 4-leaf stage. Plants were then grown in a growth chamber (E15; Conviron, Pembina, ND, USA) set at 60% relative humidity under a 12 h photoperiod (approximately 230 μmol m^-2^ s^-1^ provided by both fluorescent and incandescent bulbs) with light and dark temperatures of 22 and 20°C, respectively. Irrigation water was a 3:1 mix of two water-soluble fertilizers (15N–2.2P–12.5K and 21N–2.2P–16.6K; The Scotts Co., Marysville, OH, USA) to supply (in mg L^-1^): 200 N, 26 P, 163 K, 50 Ca, 20 Mg, 1.0 Fe, 0.5 Mn and Zn, 0.24 Cu and B, and 0.1 Mo. 76% of the nitrogen was provided as nitrate. 93% sulfuric acid (Brenntag North America, Inc., Reading, PA, USA) at 0.08 mg L^-1^ was mixed in irrigation water to maintain pH between 5.8 and 6.2. All plants were grown in the Purdue University Horticulture Plant Growth Facility (https://ag.purdue.edu/hla/Hort/Greenhouse/Pages/Default.aspx).

### Transcriptome sequencing and analysis

For RNA extraction, tissue was collected in the middle of the dark period. Four biological replicates consisting of five plants each were collected. Whole rosettes of 4-week-old plants were frozen in liquid nitrogen and stored at −80°C. Tissue was finely ground in liquid nitrogen using a mortar and pestle. Approximately 800 mg ground tissue was combined with 1 ml of TRIzol reagent (Gibco/BRL Life Technologies, Invitrogen, Carlsbad, CA, USA) and RNA was extracted according to the manufacturer’s instructions. Quality of RNA was estimated by the ratio of absorbance at 260 to 280 nm and 260 to 230 nm by spectrophotometer (DU 730; Beckman Coulter, Inc., Indianapolis, IN, USA) with both ratios between 1.8 and 2.2. RNA was reverse-transcribed into cDNA using the Poly(A) Purist protocol (Ambion, Austin, TX, USA).

The cDNA samples were fragmented into 300–500 bp molecules and sequencing libraries were constructed for the 454 GS-FLX instrument (454 Life Sciences, Roche Company, Branford, CT, USA) at the Purdue University Genomics Facility (West Lafayette, IN, USA). One sequencing run was performed (Margulies et al., 2005). Raw sequences were trimmed to remove adaptor sequences, and an initial quality trimming was performed using GS De Novo Newbler (v2.5.3; default parameters) (Part C – GS De Novo Assembler, GS Reference Mapper, SFF Tools, 2010). Trimmed reads were further trimmed using the FASTX-Toolkit (Gordon and Hannon, 2010), with a minimum quality value of 12 and minimum read length of 50 bp. Trimmed reads were aligned to the Joint Genome Institute (JGI) *E. salsugineum* genome (Phytozome v9.1: *Thellungiella halophila*: http://www.phytozome.net/thellungiella.php; Yang et al., 2013) using the splice-aware aligner GMAP v2012-11-27 with default parameters (Wu and Watanabe, 2005). Only uniquely mapped reads were used for transcript abundance estimates and single nucleotide polymorphism (SNP) calling.

SNPs were detected using mpileup from SAMtools v0.1.18 with mapping quality ≥ 15, and depth ≥ 3 (Li et al., 2009). 454 sequencing has a high error rate for detecting indels (Margulies et al., 2005), so only SNPs resulting from substitutions were retained. The two accessions are substantially inbred lines and should be homozygous at each base position. Hence, only monomorphic base positions within each accession were considered for detection of differences between the two accessions. Custom Perl scripts were used to remove SNPs 1) that were heterozygous within either accession, 2) that were not biallelic between accessions, 3) that were supported by fewer than 3 sequence reads, 4) for which the alternative allele accounts for fewer than 10% of aligned reads, and 5) that were heterozygous between the SH accession and the JGI SH reference. If more than 4 SNPs were detected within a 100 bp region using the VariantFiltration module from GATK v2.4.9 (McKenna et al., 2010), they were not included in the final SNP dataset. SNPs that had mpileup quality score 999 based SAMtools were deemed “high quality” SNPs. on are Sanger sequencing data of the YK accession available from the National Center for Biotechnology Information (NCBI) (Wong et al., 2005) was also aligned to the reference genome using SSAHA2 (Ning et al., 2001). SNPs were called using SAMtools and filtered for clustered SNPs (4 SNPs within 100 bp region) using GATK as indicated. SNPs that were not biallelic or were heterozygous within YK were removed.

Genes were identified via a reference annotation-based transcript assembly method using the Cufflinks package (Roberts et al., 2011; Trapnell et al., 2010). Reads from SH and YK were assembled separately and then merged using the *cuffmerge* command (Roberts et al., 2011; Trapnell et al., 2010). The *intersect* function within BEDTools v2.17.0 was used to identify genes not annotated in the JGI *E. salsugineum* genome (newly annotated genes). Same-strandedness was not enforced when identifying newly annotated genes because of the non-strand-specific protocol for 454 library preparation. Newly annotated genes that are unique from or overlap genes annotated by Champigny et al. (2013) but are present in the JGI reference genome were also identified using the same method (Supplementary Table 1).

**Table 1.** Number of reads, percentage of reads uniquely aligned to reference genome (%), mean read length (bp) of *E.* salsugineum Shandong (SH) and Yukon (YK) accessions.

| Accession | Total reads | Uniquely aligned (%) | Mean read length (bp) |
| --- | --- | --- | --- |
| SH | 541,494 | 95.6 | 365.8 |
| YK | 491,926 | 94.4 | 361.8 |
| Total | 1,033,420 | 95.0 | 363.8 |

The number of reads uniquely aligned to each gene was determined with htseq-count within HTSeq v0.5.4p5 (http://www-huber.embl.de/users/anders/HTSeq/doc/count.html) using *union* mode. The bioconductor package “DESeq” v.1.14.0 was used to identify genes likely to be differentially expressed between SH and YK without biological replicates (Anders and Huber, 2010). Gene expression was normalized, and the significance threshold for differential expression was based on a 0.2 false discovery rate (FDR) (Benjamini and Hochberg, 1995).

Genes were annotated by the best BLAST (Altschul et al., 1990) hit of *A. thaliana.* For *E. salsugineum* genes predicted by the JGI genome, annotation was taken from Phytozome v9.1 (Phytozome v9.1: *Thellungiella halophila*: http://www.phytozome.net/thellungiella.php Yang et al., 2013). For newly identified genes, nucleotide databases of *A. thaliana* version TAIR 10 (Lamesch et al., 2012; TAIR10: ftp://ftp.arabidopsis.org/home/tair/Sequences/blast_datasets/TAIR10_blastsets/)*, A. lyrata* (Hu et al., 2011; Phytozome v9.1: *Arabidopsis lyrata*: http://www.phytozome.net/alyrata.php), and *S. parvula* (Dassanayake et al., 2011; *Thellungiella parvula* genome: http://www.thellungiella.org/data) were built and used for similarity searching by BLASTN v2.2.28+ (Altschul et al., 1990). Genes were annotated by the best BLAST hit with the following threshold parameters: E ≤ 1^-30^; sequence identity ≥ 30%; sequence aligned ≥ 30% of query sequence.

### Quantitative real-time PCR

For quantitative real-time reverse transcription polymerase chain reaction (qRT-PCR), tissue was collected as described for 454 sequencing, RNA was extracted using the RNeasy Plant Mini Kit (QIAGEN, Valencia, CA, USA), and genomic DNA was removed using the TURBO DNA-free™ Kit (Ambion, Grand Island, NY, USA). The quality and quantity of mRNA was assessed using a NanoDrop 2000 (Thermo Fisher Scientific Inc., Wilmington, DE, USA). All samples were diluted to 200 ng μl^-1^, and 260/280 and 260/230 ratios were between 1.8 and 2.2. cDNA was synthesized using a High-Capacity cDNA Reverse Transcription Kit (Invitrogen, Grand Island, NY, USA). Primers were designed using Primer Express Software (v3.0.1). Primer specificity was then estimated by BLASTN using the *E. salsugineum* genome with all primer pairs. Primer efficiency was tested for all pairs of primers. cDNA was diluted 5 times by a 5-fold gradient, and then used as template for qRT-PCR, the threshold cycles (C_T_) were regressed against cDNA concentration (log). Slope of the regression line was estimated and the efficiency was calculated as 10^-(1/slope)^-1. For genes expressed in both accessions, primer efficiency was between 80 and 110% in both accessions. For genes that were only expressed in one accession based on RNA-seq data, primer efficiency was tested on both accessions, but only the accession with detected expression exhibited efficiency between 80 and 110%. Supplementary Table 2 contains all primer sequences except gene XLOC_004723, for which acceptable qRT-PCR primers could not be designed.

**Table 2.** Summary of transcriptome assembly of *E. salsugineum* Shandong (SH) and Yukon (YK) accessions.

|  |  |
| --- | --- |
| Number of assembled genes | 17,888 |
| Shared | 14,577 |
| SH only | 1,643 |
| YK only | 1,668 |
| Newly annotated genes | 157 |
| Shared | 122 |
| SH only | 26 |
| YK only | 9 |
| Annotated by best blast hit from *A. thaliana*, *A. lyrata* and/or *E. parvulum* | 54 |
| Newly annotated genes unique from genes annotated by Champigny et al., 2013 | 65 |
| Newly annotated genes present in genes annotated by Champigny et al., 2013 | 92 |

All qRT-PCR reactions were conducted in StepOnePlus™ Real-Time PCR Systems (Applied Biosystems, Invitrogen, Grand Island, NY, USA). Relative gene expression of target genes was quantified by the ΔC_T_ method (Livak and Schmittgen, 2001). Relative gene expression was calculated as:

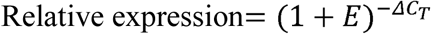

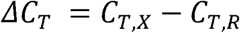

where *E* is the primer efficiency for each pair of qRT-PCR primers. *C_T,X_* and *C_T,R_* is the threshold cycle of the target gene and the reference gene *Actin2* (Thhalv10020906m.g), respectively.

### Metabolite profiling and data analysis

Two metabolite profiling experiments were conducted. One experiment was performed using the same rosette tissue used for RNA-seq analysis (see above). A second metabolite profiling experiment was performed using tissue from SH, YK, and YK x SH F_1_ plants with 3 replicates of 5 pooled plants per replicate. In both two cases, identical extraction, derivatization, and analysis methods were used. Approximately 800 mg of frozen ground tissue was incubated in methanol at 65°C in 1.75 ml tubes and centrifuged at 13,300 r min^-1^. The supernatant, containing polar molecules, was decanted into a new tube. Chloroform was added to the pellet and incubated at 37°C for 15 min to solubilize nonpolar metabolites. Samples were then dried at room temperature for about 6 h (polar) and 2 h (nonpolar) in a centrifuge at 1725 r min^-1^ and 30 μM Hg vacuum. Samples were stored at -80°C until being sent to the Metabolomics Center at the University of Illinois (http://www.biotech.uiuc.edu/metabolomics/). Samples were measured by gas chromatography/mass spectrometry (GC/MS; Agilent 6890 N/5973 MSD, Palo Alto, CA, USA) after purification and trimethylsilylation with MSTFA (*N*-methyl-*N*-trimethylsilyl-trifluoroacetamide) (Gullberg et al., 2004; Singh et al., 2011). Data was analyzed by peak identification via comparison with spectra from standards and relative concentrations of metabolites were obtained by comparison with internal standard peak area (Singh et al., 2011).

Pair-wise comparisons within each experiment were performed by two-tailed *t*-tests between SH and YK (experiments 1 and 2), and SH or YK and YK x SH F_1_ (experiment 2). In the second experiment, multiple comparisons among all three genotypes were conducted by Tukey’s studentized range test (Tukey, 1949). A *t*-test of F_1_ against the mid-parental average of SH and YK was done using the variance estimated from F_1_ hybrids. Within each experiment, genotype was treated as the only main factor. A nested analysis of variance (ANOVA) was also conducted using data from the two experiments. In the nested analysis, experiment and genotype were the two main factors. The experiment by genotype interaction was not included in the model. All identified metabolites are presented in Supplementary Table 3.

**Table 3.** Summary of single nucleotide polymorphisms (SNPs) that differentiate *E. salsugineum* Shandong and Yukon accessions.

|  |  |
| --- | --- |
| Number of genes with SNPs | 6,182 |
| Number of SNPs in genes | 14,808 |
| Number of high quality SNPs <sup>a</sup> | 4,873 |
| Number of SNPs identified by Champigny et al., 2013 | 47,317 |
| Number of SNPs present in Champigny et al., 2013 | 10,883 |
| Number of SNPs unique from Champigny et al., 2013 | 3,925 |

<sup>a</sup> Quality SNP satisfies technical and biologically relevant filtering criteria as discussed in the materials and methods.

### Pathway analysis

Pathway analysis was conducted on the annotated DEGs identified in transcriptome profiling and metabolites that differed between accessions in both metabolite experiments with the same direction. The fold differences between SH and YK was used to indicated up-regulation or down-regulation in YK compared to SH. The list of DEGs or metabolites with the fold changes were imported to MapMan software (Thimm et al., 2004). Significant pathways in which genes or metabolites were divergent from a 50/50 up/down-regulation were identified using an uncorrected Wilcoxon signed-rank (Wilcoxon, 1945).

## Results

### Novel genes and single nucleotide polymorphisms (SNPs) were identified by transcriptome sequencing

Whole rosettes of 4-week-old YK and SH plants grown in a 12h:12h light dark cycle were harvested in the middle of the dark period. Libraries of cDNA isolated from these rosettes were sequenced by pyrosequencing. These reads were aligned to the *E. salsugineum* reference genome, which was constructed using DNA from the SH accession (Yang et al., 2013). More than 1 million cDNA sequence reads, 95% of which aligned to the reference genome, were used for a reference-directed assembly of the transcriptome (Table 1), identifying 17,888 expressed genes (Table 2 and Supplementary Table 1). Of these, 65 genes were novel and not predicted in the reference genome (Yang et al., 2013) nor detected in a previous transcriptome analysis conducted by Champigny et al. (2013). Only 20 of these 65 genes have annotated orthologs in the related species *A. thaliana*, *A. lyrata*, and/or *Schrenkiella parvula* (Supplementary Table 1). Presence-absence variation (PAV), defined as zero reads aligned to one of the two parents, was observed for 18.5% of the detected expressed genes with roughly equal numbers of genes detected only in SH or YK (Table 2).

The transcript assemblies were processed to detect SNPs between SH and YK. 42% of shared genes contained a total of 14,808 SNPs, of which 4,873 were deemed “high quality” (Table 3; Supplementary Table 4). We compared our results with the SNP detection results of Champigny et al. (2013). Of the SNPs detected in our experiment, 73% (10,883 positions) and 79% (3861 positions) of the low and high stringency SNPs, respectively, were also identified by Champigny et al. (2013). We also compared our SNPs to available Sanger sequencing of cDNA clones from the YK accession (Wong et al., 2005) and identified 468 putative SNPs with reference to the SH reference genome. Of these, 441 have corresponding sequence data in the YK transcriptome we assembled and 88% (388 SNPs) had the same sequence variation in our assembly and the YK Sanger sequencing data (Supplementary Table 4).

**Table 4.**
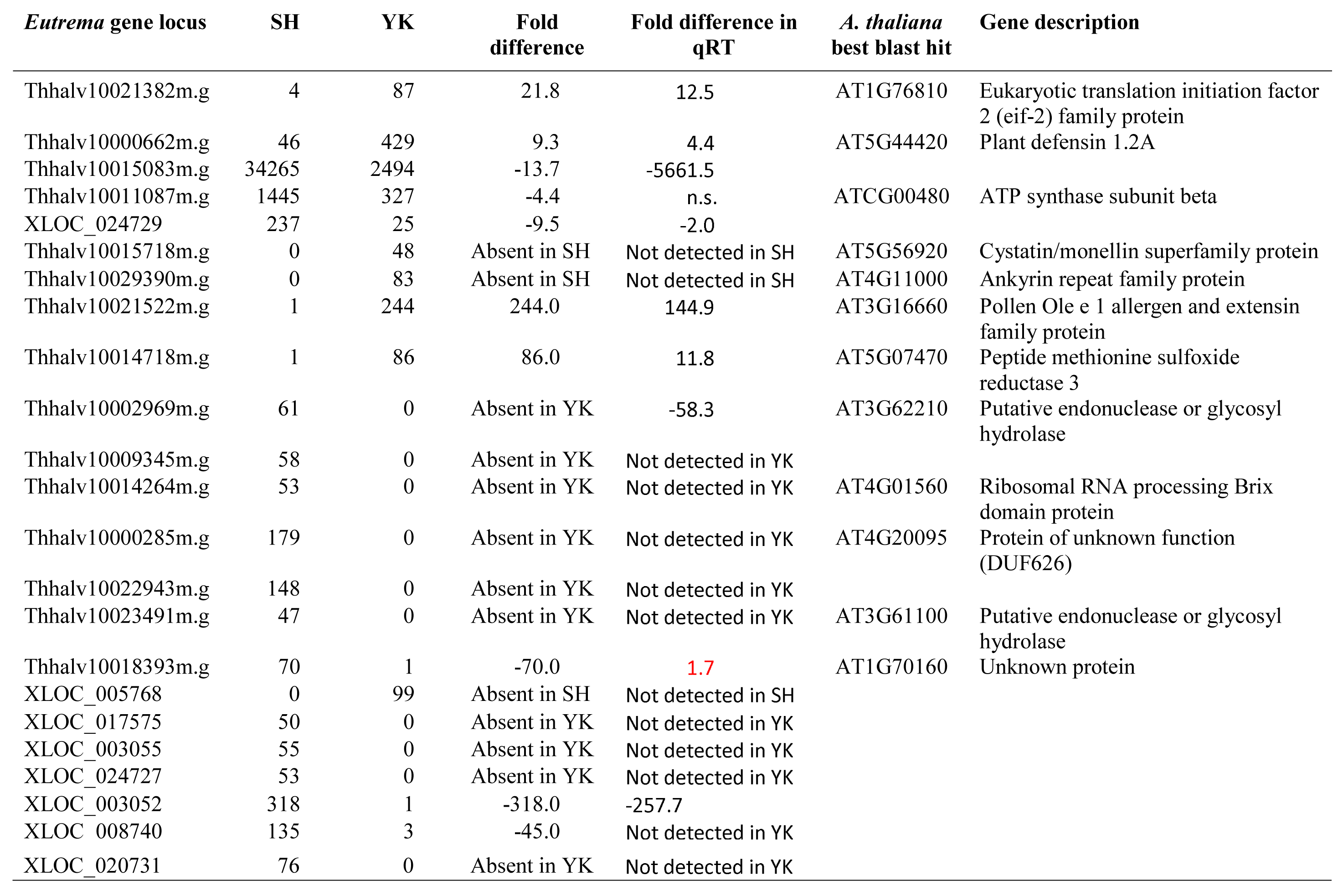
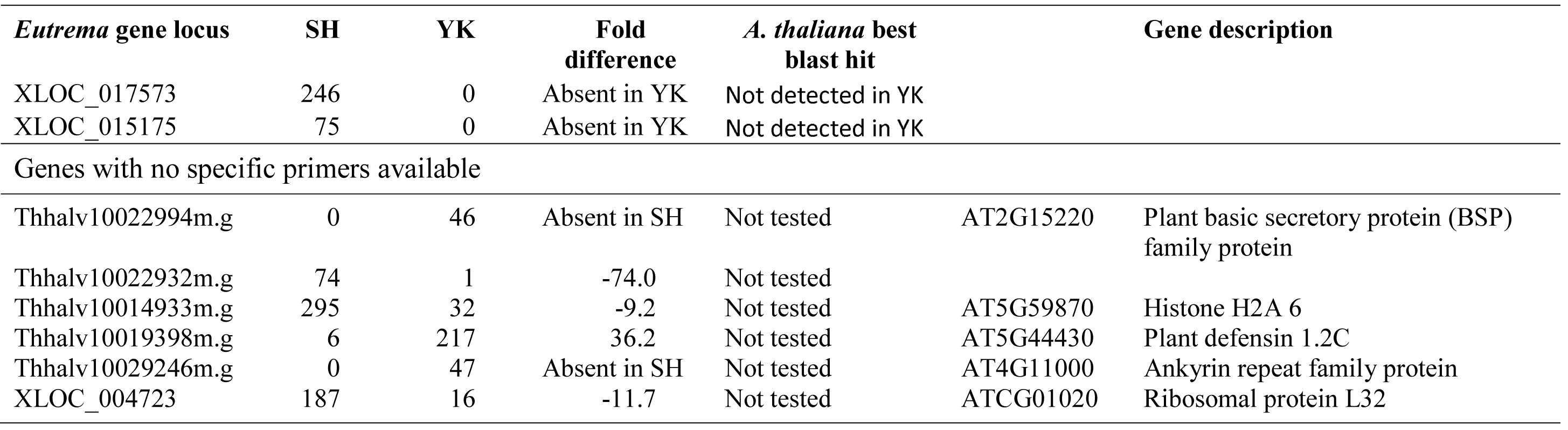
Differently expressed genes (DEGs) of *E. salsugineum* Shandong (SH) and Yukon (YK) accessions.

### Transcriptome profiling and qRT-PCR identify differentially expressed genes (DEGs) between SH and YK

To assess gene expression differences between SH and YK, we determined the number of reads aligned to each gene. Less than 0.2% (thirty-one genes) of the expressed genes were identified as candidate DEGs between SH and YK (Table 4). Sixteen of these thirty-one candidates do not have homologous genes in *A. thaliana*, *A. lyrata*, or *S. parvula* and twenty have been annotated in the reference genome (Table 4). Of those with homologs in one of these species, none have been associated with previously reported trait differences in SH and YK.

To confirm DEGs based on RNA-seq, we designed quantitative real-time reverse transcription polymerase chain reaction (qRT-PCR) primers that matched unique positions in the reference genome based on BLAST analysis (Altschul et al., 1990). Unique primers could not be designed for five genes (Thhalv10022994m.g, Thhalv10022932m.g, Thhalv10014933m.g, Thhalv10019398m.g, and Thhalv10029246m.g) due to paralogs with high sequence similarity and no acceptable primer pair was identified for XLOC_004723. In total, qRT-PCR data confirmed our RNA-seq data for 23 of 25 genes. For the 5 DEGs that had no close paralogs and at least 4 reads in both SH and YK, expression differences based on RNA-seq were confirmed by qRT-PCR in 4 of 5 genes (Table 4). Among the 20 genes that had fewer than 4 reads in either SH or YK, expression differences by qRT-PCR was consistent with RNA-seq data in 19 (Table 4). In addition, when the low accession had fewer than 4 reads there was no amplification in 16 of 20 cases (Table 4).

### Metabolite profiling reveals higher accumulation of fatty acids and amino acids in YK and enhanced soluble carbohydrate accumulation in SH

To identify potential metabolites and metabolic pathways that contribute to phenotypic and physiological differences between SH and YK, metabolite profiling was conducted in two independent experiments. In one experiment, metabolite concentrations in F_1_ plants of a YK x SH cross were also determined.

Concentrations of free fatty acids and long-chain fatty acid derivatives were higher in YK than SH (Table 5 and Supplementary Table 3). The concentration of ferulic acid was also greater in YK than SH, indicating a potential for greater suberin and/or cutin accumulation in the YK accession.

**Table 5.** Metabolite profiling differences between *E. salsugineum* Shandong (SH) and Yukon (YK) accessions. Metabolites for which tissue concentrations were different between *E. salsugineum* Shandong (SH) and Yukon (YK) accessions in one or two experiments and/or different based on a nested analysis of the two experiments.

| Metabolite | Experiment 1 |  | Experiment 2 |  | P-value |  |  |
| --- | --- | --- | --- | --- | --- | --- | --- |
|  | SH | YK | SH | YK | E1 | E2 | N |
| Carbohydrates |  |  |  |  |  |  |  |
| 2-keto-gluconic acid | 1012.0 | 526.4 | 352.9 | 470.8 | n.s. | * | n.s. |
| 2PGA | 2.1 | 0.0 |  |  | ** |  |  |
| α-ketoglutaric acid | 83.2 | 41.3 | 66.5 | 61.2 | * | n.s. | * |
| Arabitol | 45.4 | 60.9 | 12.9 | 19.5 | n.s. | * | n.s. |
| Ascorbic acid | 19.6 | 11.8 | 25.0 | 10.4 | n.s. | ** | ** |
| Citric acid | 6722.6 | 17762.7 | 14347.0 | 30526.0 | n.s. | ** | ** |
| Digalactosylglycerol | 2007.8 | 618.8 | 344.5 | 82.9 | n.s. | *** | n.s. |
| Disaccharide | 9941.3 | 8651.1 | 10713.7 | 3862.7 | n.s. | *** | ** |
| Erythritol | 27.6 | 39.1 | 22.1 | 34.3 | n.s. | * | * |
| Erythrose |  |  | 976.8 | 591.3 |  | * |  |
| Fructose | 65604.0 | 9500.3 | 14987.4 | 1284.4 | *** | *** | *** |
| Fumaric acid | 232.3 | 429.0 | 111.5 | 263.1 | n.s. | *** | ** |
| Galactaric acid |  |  | 3.4 | 8.5 |  | * |  |
| Galactose | 7730.3 | 4432.0 | 32.0 | 88.7 | * | *** | ** |
| Gentiobiose |  |  | 27.3 | 20.4 |  | * |  |
| Glucaric acid | 3.7 | 3.0 | 16.1 | 10.4 | n.s. | * | n.s. |
| Glucose | 41843.5 | 28552.0 | 16547.4 | 6251.4 | * | *** | *** |
| Glycerol-3-p | 82.4 | 30.8 | 674.0 | 362.7 | * | *** | *** |
| Isomaltose | 75.1 | 0.0 |  |  | ** |  |  |
| Itaconic acid | 142.0 | 32.2 |  |  | * |  |  |
| Maltose | 1119.8 | 678.6 | 1081.4 | 846.2 | * | * | *** |
| Melibiose | 0.0 | 5.4 |  |  | *** |  |  |
| Methylmaleic acid |  |  | 16.7 | 7.1 |  | * |  |
| Pyruvic acid | 42.4 | 38.6 | 59.8 | 44.0 | n.s. | ** | n.s. |
| Raffinose | 477.5 | 322.5 | 218.9 | 152.7 | * | n.s. | ** |

**Table 5 Metabolite profiling differences between *E.**
| Metabolite | Experiment 1 |  | Experiment 2 |  | P-value |  |  |
| --- | --- | --- | --- | --- | --- | --- | --- |
|  | SH | YK | SH | YK | E1 | E2 | N |
| Sedoheptulose | 853.9 | 782.4 | 452.2 | 312.6 | n.s. | ** | n.s. |
| Sorbitol | 17.0 | 39.5 |  |  | * |  |  |
| Turanose |  |  | 13688.2 | 7035.2 |  | ** |  |
| Xylitol | 18.6 | 11.5 | 13.7 | 20.4 | n.s. | * | n.s. |
| Xylose | 165.8 | 81.1 | 64.6 | 49.3 | * | n.s. | * |
| Fatty acids |  |  |  |  |  |  |  |
| 13-eicosenoic acid |  |  | 5.7 | 7.6 |  | * |  |
| 7-hexadecenoic acid |  |  | 132.2 | 27.0 |  | * |  |
| 9,12-octadecadienoic acid |  |  | 6032.7 | 7434.6 |  | * |  |
| Docosanoic acid | 50.5 | 241.3 | 176.5 | 1224.1 | ** | *** | *** |
| Hexacosanoic acid | 13.2 | 28.9 | 0.0 | 20.9 | * | *** | *** |
| Hexadecanol | 0.0 | 12.2 |  |  | *** |  |  |
| Octacosanol | 9.5 | 1.9 |  |  | ** |  |  |
| Tetracosanoic acid | 1206.3 | 5001.7 | 8103.8 | 16232.9 | * | * | ** |
| Tetracosanol | 2.2 | 0.0 | 2.5 | 4.5 | *** | ** | n.s. |
| Tricosanoic acid | 0.0 | 12.6 | 133.6 | 150.7 | ** | n.s. | n.s. |
| Tritriacontanol |  |  | 47.1 | 37.6 |  | * |  |
| Amino acids |  |  |  |  |  |  |  |
| Alanine |  |  | 2.7 | 6.8 |  | ** |  |
| Aspartic acid |  |  | 928.9 | 2095.8 |  | * |  |
| Glycine |  |  | 13.2 | 57.0 |  | * |  |
| N-acetylglutamic acid |  |  | 158.0 | 1221.7 |  | * |  |
| Serine |  |  | 216.4 | 818.2 |  | * |  |
| Threonine | 0.0 | 3.8 | 311.1 | 383.3 | *** | * | * |

**Table 5 Metabolite profiling differences between *E.**
| Metabolite | Experiment 1 |  | Experiment 2 |  | P-value |  |  |
| --- | --- | --- | --- | --- | --- | --- | --- |
|  | SH | YK | SH | YK | E1 | E2 | N |
| Valine | 0.0 | 10.3 | 104.9 | 92.8 | ** | n.s. | n.s. |
| Others |  |  |  |  |  |  |  |
| 1,2,3-hydroxybutane | 4.1 | 0.0 |  |  | * |  |  |
| 2,3-dihydroxybutanedioic acid | 11.8 | 27.0 | 5.8 | 9.6 | * | * | ** |
| 2,4,6-tri-tert.-butylbenzenethiol | 42.5 | 44.3 | 157.3 | 185.3 | n.s. | * | n.s. |
| 2-aminoethylphosphate |  |  | 0.0 | 44.5 |  | * |  |
| 2-desoxy-pentos-3-ulose | 33.1 | 15.3 |  |  | *** |  |  |
| 2-hydroxybutanoic acid |  |  | 11.3 | 0.0 |  | * |  |
| 2-o-glycerol-galactopyranoside | 2362.0 | 398.0 | 181.6 | 178.0 | *** | n.s. | * |
| 2-oxo-3-hydroxypropanoic acid |  |  | 17.3 | 75.7 |  | ** |  |
| 3-methylthiopropylamine |  |  | 0.0 | 5.4 |  | * |  |
| 5-o-coumaroyl-d-quinic acid |  |  | 38.5 | 15.6 |  | * |  |
| Aconitic acid | 202.7 | 103.5 | 12.7 | 37.5 | * | * | n.s. |
| Tocopherol | 7.8 | 4.0 |  |  | * |  |  |
| Chlorogenic acid |  |  | 710.6 | 332.2 |  | * |  |
| Erythronic acid-1,4-lactone |  |  | 24.2 | 38.0 |  | * |  |
| Ferulic acid | 11.3 | 30.4 | 41.8 | 61.6 | n.s. | ** | ** |
| Hydroxylamine |  |  | 39.2 | 27.1 |  | ** |  |
| Ketomalonic acid |  |  | 0.0 | 3.1 |  | * |  |
| Malonic acid | 59.7 | 89.5 | 0.0 | 2.6 | n.s. | ** | n.s. |
| P-hydroxyacetophenon | 0.0 | 2.8 |  |  | *** |  |  |
| Protocatechuic acid |  |  | 5.9 | 2.1 |  | * |  |

**Table 5 Metabolite profiling differences between *E.**
| Metabolite | Experiment 1 |  | Experiment 2 |  | P-value |  |  |
| --- | --- | --- | --- | --- | --- | --- | --- |
|  | SH | YK | SH | YK | E1 | E2 | N |
| Putrescine |  |  | 0.0 | 22.4 |  | ** |  |
| Stigmasterol | 40.6 | 52.0 | 29.5 | 9.1 | n.s. | ** | n.s. |
| Vanillin | 0.0 | 1.8 |  |  | *** |  |  |

Data are means (n=4). Empty indicates absence or undetected metabolites. Genotypes are significant different at P- value < 0.05 (\*), 0.01 (\*\*), and 0.001 (\*\*\*) based on two-tailed t test in Experiment 1 (E1), Experiment 2 (E2), or nested (N) analysis of two experiments combined together. Carbohydrates indicate sugar-related compounds and derivatives in citric acid cycle. Fatty acids indicate fatty acids and their derivatives. Others indicate all other compounds not belong to carbohydrates or fatty acids, or amino acids.

The products of starch degradation were more abundant in SH than YK in both profiling experiments (Table 5). Maltose and glucose, primary products of starch degradation, were elevated in SH along with fructose, glycerol-3-phosphate, raffinose, and an unresolved disaccharide. This suggested more active 6-carbon metabolite catabolism via glycolysis in SH compared to YK during the night (Table 5 and Supplementary Table 3). SH accumulated higher concentrations of the disaccharides isomaltose and gentiobiose while YK accumulated a very small amount of melibiose in one of the two screens. Tricarboxylic acid (TCA) cycle intermediates, on the other hand, were not consistently different between the accessions. Citric acid and fumaric acid were accumulated at higher levels in YK while alpha-ketoglutaric acid was greater in SH (Table 5 and Supplementary Table 3). Ascorbic acid was higher in SH but the ascorbic acid degradation product tartaric acid (2,3 dihydroxybutanedioc acid, Table 5 and Supplementary Table 3) was higher in YK.

All amino acids that differed between SH and YK were higher in YK (Table 5 and Supplementary Table 3). These included alanine, glycine, serine, threonine, and valine. Of these, only threonine was differentially accumulated in both profiling experiments. The amino acids alanine, glycine, and serine were only detected in one of the two experiments. Valine was detected in both experiments, but the difference between SH and YK was only statistically significant in the first experiment. Concentrations of the other detected amino acids (isoleucine, leucine, and proline) were not different in the two accessions.

Pathway analyses were conducted to on both transcriptome (Table 4) and metabolome data (Table 5) to identify differences between SH and YK during the dark period and any correlation between transcriptome and metabolome analyses of the two accessions. The metabolite pathway analysis identified two functional categories: major carbon degradation (down-regulated in YK) and amino acid synthesis (up-regulated in YK). Glucose, maltose, and isomaltose are within the major carbon degradation pathway, while aspartic acid, alanine, N-acetylglutamic acid, threonine, serine, and glycine are within the amino acid synthesis pathway. However, of the 19 DEGs annotated in the reference genome, one unknown functional category including nine DEGs (Thhalv10015083m.g, Thhalv10002969m.g, Thhalv10023491m.g, Thhalv10021522m.g, Thhalv10018393m.g, Thhalv10009345m.g, Thhalv10015718m.g, Thhalv10022932m.g, and Thhalv10022943m.g) and one category related to cell organization of two DEGs (Thhalv10029390m.g and Thhalv10029346m.g) were identified. Hence, our analyses did not show any correlation between the *E. salsugineum* transcriptome and metabolome.

### Metabolomic phenotypes in the F_1_ hybrid exhibit transgressive heterosis

To better understand the genetic basis for observed differences in metabolite concentrations between SH and YK, we measured metabolites in F_1_ hybrids. Concentrations of all measured metabolites in both experiments are presented in Supplementary Table 3. Of the 144 metabolites measured in the experiment comparing the two parents and hybrids, 56% of all metabolites were lower in hybrids than the predicted mid-parent value. This included 65% of the fatty acids and 49% of carbohydrates detected (Supplementary Table 3). We performed a two-way contingency test to determine if an observed difference in the accumulation of a metabolite was predictive of heterosis for that metabolite (Supplementary Table 5). We found that metabolites with accumulation differences between the parents were neither more nor less likely to exhibit accumulation differences between the F_1_ and the mid-parental values (Supplementary Table 5).

Hybridization can result in transgressive heterosis in which phenotypic values for the hybrids fall outside of the range of parental values. Of the 144 metabolites measured, transgressive heterosis was observed for 28. Of these, 4 metabolites were not observed in one of the two parents and 24 were detected in both parents and the hybrids (Figure 1). Of the 24 detected in all three genotypes, the transgressive heterosis was more likely to be negative (Figure 1; *P*-value ≤ 0.0001 based on Binomial exact test) and more frequently affected metabolites that did not differ in concentration between the parents (Supplementary Table 5; *P*-value ≤ 0.05 based on χ^2^ test). Thus, heterosis for the metabolome was manifested by a decrease in metabolite pool sizes in hybrids and was not preferentially associated with metabolites that contributed to the variation between the two parents. This is consistent with our observation of decreased availability of primary metabolites in the faster growing YK (Table 5 and Supplementary Table 3). We propose that a metabolic consequence of enhanced growth is a reduction in pool sizes of primary metabolites and greater resource utilization for anabolic metabolism.

**Figure 1.**
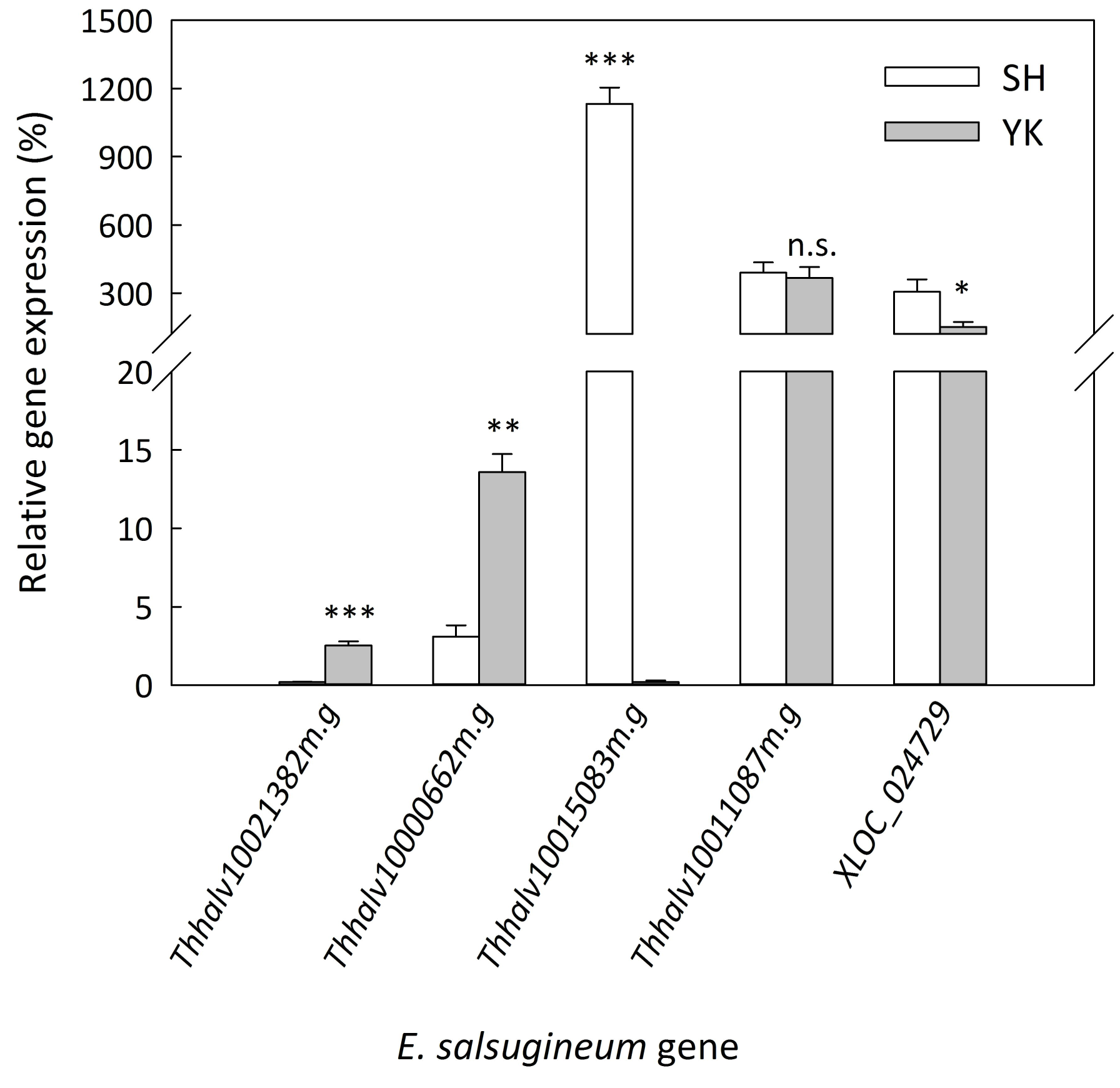
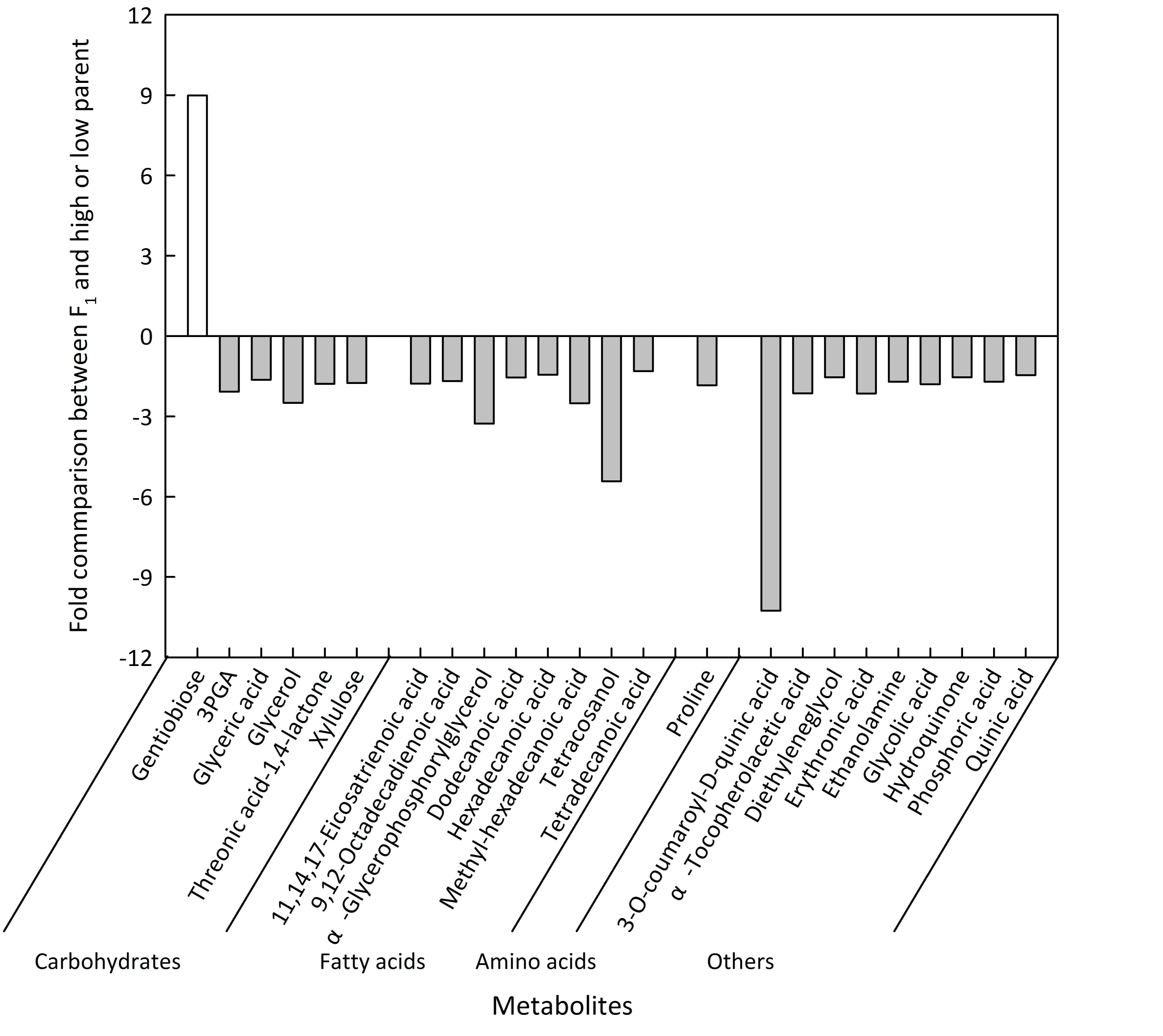
Heterosis for metabolite concentrations in F_1_ hybrids. Metabolites in F_1_ hybrids that were higher than the high parent or lower than the low parent at *P*-value≤0.05 based on two-tailed *t*-test are shown. The ratio is calculated as F_1_/high parent when F_1_ had a higher concentration than the high parent and low parent/F_1_ when F_1_ had a lower concentration than the low parent.

## Discussion

In this study, we profiled both transcript and metabolite accumulation to identify genetic and biochemical variation during the dark period in two *E. salsugineum* accessions: SH and YK. We annotated novel genes present in the reference genome and identified DEGs between SH and YK in the middle of the dark period. We found that YK accumulates more fatty acids than SH, while SH accumulates sugars at higher concentrations. Although the transcriptomic and metabolic profiling results do not offer links to each other, they do offer insight into genetic and physiological differences between these accessions. Furthermore, we identified additional SNPs, and provide validation of previously described SNPs, varying between these accessions that can be utilized for future research.

### Validity of identified SNPs

Based on the predicted transcriptome size (Yang et al., 2013), the SNP density from our analysis is 1 SNP per 10 kb of transcribed sequence. 14,887 SNPs identified from our transcriptome sequencing data were also present in Champigny et al. (2013). However, our SNP cluster filter, which removes neighboring SNPs to account for misaligned reads at insertion-deletion polymorphisms, removed 3,604 overlapping SNPs. The removal of this set of SNPs, likely enriched for false positives, contributed to the lower SNP number detected in our study. We also acknowledge that this contributed to our false negative rate. Of the 388 SNPs that matched between our analyses of pyrosequencing data and the Sanger data, 99 were removed by our procedure. We provide a compact file containing the subset of “high-quality” (see Materials and Methods) SNPs in the supplementary data. These have a SNP density of 1 SNP per 25 kb of transcript (Supplementary Table 4). This set of validated SNPs can be used for QTL mapping and fine-mapping studies (Matsuda et al., 2015; Trick et al., 2012; Wu et al., 2016).

### Identification of genes expressed in the dark period in *E. salsugineum*

We provided expression data supporting 63% and 67% of the predicted genes in the two published reference genomes (Wu et al., 2012; Yang et al., 2013). This is likely an underestimate of the expressed genes in this species because we sampled only leaf tissue and only at night (Birnbaum et al., 2003; Czechowski et al., 2004; Doherty and Kay, 2010; Huang et al., 2012b). We identified 66% of the genes identified in another transcriptome analysis of *E. salsugineum* (Champigny et al., 2013). Despite our study relying on lower coverage (4X vs. 8X), and quantifying expression of genes only expressed in leaves at night, we annotated 65 genes not predicted in the reference genome or identified in the previous transcriptome characterization (Table 2 and Supplementary Table 1). Novel genes identified in this transcriptome study but not that of Champigny et al. (2013) are likely either only expressed at night or otherwise not expressed under the conditions of the previous study which included a 21 hr day and low dark period temperatures.

More than 97% of the genes identified in our study are homologous to genes in *A. thaliana* (Supplementary Table 1), consistent with the relatively close phylogenetic association of the two species (Al-shehbaz et al., 1999). Of the 65 newly-annotated genes in this study, 47 of them do not have homologs in *A. thaliana*. One of these novel genes has been annotated in *A. lyrata* and two have been annotated in *S. parvula* (Supplementary Table 1). Therefore, we may have identified genes that are unique to this extremophile species (Wu et al., 2012).

More than 80% of the expressed genes identified in this study were detected in both SH and YK (Table 2). This indicates a conserved transcriptome between these two accessions and is consistent with previous *E. salsugineum* transcriptome comparisons (Champigny et al., 2013; Lee et al., 2013). PAVs, as scored by read count in the transcript profiling experiment, accounted for 19 and 12% of all genes in this experiment and Champigny et al. (2013), respectively. Only 13% of these were consistently detected as PAV over the two datasets. These genes are likely “true” PAV genes (Supplementary Table 1). Other genes detected in only one of the replicates were likely due to low mRNA abundance. PAV structural variation has been observed in *A. thaliana* (Bush et al., 2014), maize (Springer et al., 2009; Swanson-Wagner et al., 2010), and soybean (Haun et al., 2011). PAV genes are not typically essential (Bush et al., 2014) and may have minor effects on plant fitness (Swanson-Wagner et al., 2010). However, genes present in only one accession could contribute to the adaptation to specific selective constraints (Bush et al., 2014) and variation in quantitative traits (Swanson-Wagner et al., 2010). The observed phenotypic variation in growth rate and metabolism could be due to PAV, although no identified PAVs have been linked to trait differences in *Eutrema*. Further study utilizing molecular genetics to address the causes and consequences of natural variation in this species is needed to link these data types.

### DEGs and constitutive response to abiotic and biotic stresses

The low number of candidate DEGs between the two accessions of *E. salsugineum* used in this study was similar to other studies. 55% of our these had no reads in one of the two accessions, which could be due to a lack of sufficient depth to detect very low expression. Champigny et al. (2013) identified 381 DEGs that were present in our transcriptome but not identified as DEGs in our data. RNA-seq experiments are often underpowered, and the lack of overlap with the previous study may be primarily an issue of low read depth and replicate numbers. Consistent with this expectation, 58% of the 381 DEGs identified previously have low expression (<4 reads; Supplementary Table 1). In addition, 41% of the DEGs from Champigny et al. (2013) exhibit lower expression during the dark period in *A. thaliana* (Mockler et al., 2007), which may explain some of the lack of overlap in DEGs in the two studies. We identified 17 DEGs that were not identified by Champigny et al. (2013), possibly because they are only differentially expressed during the dark period or under our experimental conditions.

It appears that the relative abundance of most transcripts are similar in SH and YK, since only 0.2 and 1.9% of all genes were DEGs in our study and Champigny et al. (2013), respectively. Fourteen of the 31 DEGs identified in our study were also identified by Champigny et al. (2013) and the expression patterns were the same in both studies. This suggests that differential expression for these genes between SH and YK is consistent over light and dark periods and the two growth conditions.

Without exposure to abiotic or biotic stresses, YK expressed several stress-responsive genes (Table 4). This was also noted by Champigny et al. (2013). Two plant defensin genes within the same family were highly expressed in YK (Table 4). In *A. thaliana,* the plant defensin type 1 family (*PDF1*) is comprised of seven genes (Shahzad et al., 2013) with highly conserved sequences and identical mature peptides (Thomma et al., 2002). *AtPDF1* genes are induced by pathogens, non-host pathogens, methyl jasmonate (MeJA), and ethylene (ET) (De Coninck et al., 2010; Manners et al., 1998; Penninckx et al., 1996; Zimmerli et al., 2004). Also, expression of *AtPDF1s* in yeast results in zinc tolerance (Shahzad et al., 2013). Eight gene models in *E. salsugineum* are homologous to the *A. thaliana* plant defensin family. Of these, one gene is annotated as the likely ortholog of *AtPDF1.4*, whereas five are annotated as most closely related to *AtPDF1.2A*, and two are annotated as *AtPDF1.2C*. Higher expression of *PDF1.2* and *PDF1.4* was observed in the YK accession, however, paralogous *PDF1* genes are expressed in SH (Supplementary Table 1).

The transcript abundance of a gene encoding *E. salsugineum* peptide methionine sulfoxide reductase 3 (*PMSR3*) was higher in YK than SH (Table 4). There are five orthologous *PMSR* genes in *A. thaliana*: *PMSR1* to *5* (Rouhier et al., 2006) that are also found in *E. salsugineum*. The expression of *PMSR3* is induced by arsenate (Paulose et al., 2010), suggesting the ability to respond to plant abiotic stresses. No function in tolerance or resistance has been established for this paralog in *A. thaliana*. However, knock-out of either *PMSR2* (Bechtold et al., 2004) or *PMSR4* (Romero et al., 2004) results in decreased oxidative stress tolerance and overexpression of either gene increases stress tolerance in *A. thaliana*. Expression of *PMSR4* (but not *PMSR1*, *PMSR2*, or *PMSR3*) was induced in response to UV and AgNO_3_ in *E. salsuginea* SH (Mucha et al., 2015). It is plausible that the overexpression of *PMSR3* by YK could provide greater oxidative stress tolerance in this accession.

### *E. salsugineum* accessions SH and YK differ in carbon metabolism

In two metabolic profiling experiments, 125 and 144 metabolites were detected. Differences were identified between SH and YK for the 85 metabolites detected in both experiments (Table 5 and Supplementary Table 3). The derivatization method we utilized has been widely used to detect sugars (Gullberg et al., 2004), but is less accurate for identifying and quantifying amino acids (Kaspar et al., 2009). As a result, despite the fact that Eutrema accumulates higher concentrations of some amino acids than Arabidopsis (Eshel et al., 2017), many amino acids were not detected in our analyses.

Higher concentrations of fatty acids and fatty acid derivatives were measured in YK, including several previously identified as structural components of membrane lipids, cuticle components, and wall-resident suberin (Table 5 and Supplementary Table 3). Fatty acids contain more energy than carbohydrates when used as storage compounds and can act as an efficient storage form of reduced carbon (Taiz and Zeiger, 2010). High production of fatty acids is typical of rapidly growing tissues (Ohlrogge and Jaworski, 1997; Qin et al., 2007). Very-long-chain fatty acids (VLCFAs; C20:0 to C30:0) play an important role in cell elongation and expansion (Qin et al., 2007). Several VLCFAs, including docosanoic, hexacosanoic, pentacosanoic, tetracosanoic, and tricosanoic acids were more abundant in YK than SH (Table 5), consistent with our measurements of higher growth rates of YK as compared to SH in our growth conditions (manuscript in preparation). The VLCFA tetracosanoic acid, which plays an important role in root cell growth and expansion (Qin et al., 2007), was accumulated at a higher concentration in YK (Table 5 and Supplementary Table 3). In addition to carbon storage, lipids are important components of membranes and the leaf cuticle (Lynch and Dunn, 2004; Tresch et al., 2012; Zäuner et al., 2010). Our results are in agreement with Xu et al. (Xu et al., 2014), who measured a greater accumulation of C22 and C24 fatty acids in the epicuticular wax of YK over SH.

Overall, SH tissues had higher concentrations of sugars than YK. These measurements of nighttime sugar concentrations were similar to previous results obtained for fructose, glucose, and raffinose in leaves of *E. salsugineum* harvested during the day (Lee et al., 2012; Eshel et al., 2017). The products of starch breakdown, including maltose and glucose, were more abundant in SH (Table 5 and Supplementary Table 3), suggesting a higher rate of starch metabolism in the lower-biomass SH accession This was consistent with the strong negative correlation between starch content and biomass observed in *A. thaliana* accessions (Sulpice et al., 2009).

A study of the correlation between specific metabolites and biomass accumulation in *A. thaliana* revealed twenty-three metabolites that were correlated with biomass (Meyer et al., 2007). We detected fourteen of these twenty-three metabolites (Table 5 and Supplementary Table 3). Of these, five differed between SH and YK. The concentrations of ascorbic acid, glycerol-3-phosphate, and raffinose were negatively correlated and putrescine was positively correlated with biomass in *Arabidopsis* (Meyer et al., 2007). The levels of these metabolites also corresponded to the differences in biomass between SH and YK, indicating that the relationships between metabolites and biomass found in *A. thaliana* were consistent in *E. salsugineum* (Table 5 and Supplementary Table 3). This suggests that, although stress tolerance is vastly different between these two species (Amtmann, 2009; Griffith et al., 2007; Lee et al., 2012; Volkov et al., 2003), the metabolic markers for biomass accumulation may be similar.

### Increased utilization rate as a hypothesis for metabolome heterosis

More than 58% of the metabolites in F_1_ plants were different from the predicted mid-parent concentration (Supplementary Table 3), indicating a non-additive effect of hybridity on the majority of the metabolome. The lower concentration of fructose and glucose in F_1_ hybrids suggests high rates of starch and sugar depletion to support rapid growth (Lisec et al., 2011). Transgressive heterosis was more commonly observed for metabolites that were not different between the two parents (Supplementary Table 5). This suggests that allelic variation affecting differential metabolite accumulation in the parents is not responsible for the observed heterosis in the F_1_ metabolome. Although it is surprising that differences between the parents were not predictive of a metabolite association with heterosis, it may be that the metabolomic consequences of heterosis derived from secondary effects of an increased growth rate in F_1_ hybrids, rather than a causative relationship between growth rate and specific metabolites or metabolite diversity. Differences in biomass polymers and metabolites involved in anabolic growth exhibited reduced pool sizes in the more rapidly growing YK as compared to SH (Table 5 and Supplementary Table 3), as well as in the very rapidly growing F_1_ plants as compared to the parents (Figure 1 and Supplementary Table 3). Consistent with the hypothesis that utilization rate drives the heterotic effects on metabolite pool sizes, the transgressive effect overwhelmingly resulted in lower concentrations of metabolites in the hybrids (Figure 1 and Supplementary Table 3 and 5). This hypothesis regarding the cause of metabolic heterosis may be a general phenomenon in plants. Indeed, the same associations have been previously observed in maize, in which largely negative overdominance for metabolites was found in the heterotic B73 x Mo17 hybrids (Lisec et al., 2011).

## Conclusions

Our study contributes to the annotation of the *E. salsugineum* genome and provides evidence of transcriptional and metabolic differences between the SH and YK accessions. Very few differences in gene expression were detected in the middle of the dark period between these two accessions, but YK has constitutively higher expression of several plant systematic defense genes. The high-quality SNPs identified in this study can be used with previously identified SNPs to map traits that differ in these accessions, such as tolerance to various stresses. There is evidence for contrasting carbon metabolism in these two accessions, which correlates with observed growth differences. Furthermore, metabolite profiling of the accessions and F_1_ hybrids supports the notion that the concentrations of key metabolites are correlated with growth rate, including the increased growth rate caused by heterosis.

Our hypothesis was that combined transcriptome and metabolome profiling of two contrasting *E. salsugineum* accessions might elucidate the pathway(s) related to the phenotypic differences between these contrasting accessions. The difference carbon metabolism identified in the metabolome profiling provides insights for growth differences between SH and YK. However, none of the 19 DEGs that have been annotated in the reference genome are related to the observed metabolic differences. There are two plausible explanations: 1) the additional 11 DEGs that are currently unannotated in the reference genome could provide additional evidence for the link between metabolome and transcriptome, or 2) by increasing the number of replicates in transcriptome study, more DEGs will be identified to support further pathway identification.

## Availability of supporting data

All the raw data supporting the results of this article have been deposited at (Edgar et al., 2002) are accessible through GEO Series accession GSE GSE67745 at (http://www.ncbi.nlm.nih.gov/geo/query/acc.cgi?acc=GSE67745).

## Abbreviations

ABA: Abscisic acid
DEGs: Differentially expressed genes
ESTs: Expressed sequence tags
ET: Ethylene
FDR: False discovery rate
GC/MS: Gas chromatography mass spectrometry
JA: Jasmonic acid
MeJA: Methyl jasmonate
PAV: Presence-absence variation
PDF: Plant defensin gene
PMSR: Peptide methionine sulfoxide reductase
QTL: Quantitative trait loci
SH: *E. salsugineum* Shandong accession
SNP: Single nucleotide polymorphism
VLCFA: Very long-chain fatty acids
YK: *E. salsugineum* Yukon accession.

## Authors’ contributions

JY and MJG performed the experiments; JY, BPD, and MVM designed the experiments; JY, and BPD analyzed the data; JY, BPD, and MVM wrote the manuscript.

### Acknowledgements

Partial support to JY was provided by the U.S. Department of Agriculture, National Institute of Food and Agriculture – Agriculture and Food Research Initiative Grant no. 2010-85117-20607, the China Scholarship Council, and the Purdue Department of Horticulture and Landscape Architecture. We thank Dr. Scott J. Emrich and Dr. Allison Regier (current address: Genome Institute, Washington University) in the Department of Computer Science and Engineering at University of Notre Dame and Dr. Michael Gribskov in the Department of Biological Sciences for assistance and discussion of RNA-seq data analysis. We thank Dr. Chan yul Yoo (current address: Biology Department, Duke University) for the assistance of RNA-seq sample collection and preparation. We thank Rob Eddy and Dan Hahn for assistance with plant growth, Dr. Phillip San Miguel and the Purdue University Genomics core for RNAseq data production and transcriptome analysis, Dr. Alexander Ulanov and Dr. Zhong (Lucas) Li at the Metabolomics Center at the University of Illinois for assistance with the metabolomic data, and Prof. David Rhodes for discussions regarding the metabolite data.

## Supplementary data

**Supplementary Table 1. All genes identified in *E. salsugineum* Shandong (SH) and Yukon (YK) accessions.** Information for all identified genes includes a gene identification number (Gene_id) as annotated in the JGI reference genome (Thhalv) or newly annotated (XLOC), the number of raw reads in Shandong (SH) or Yukon (YK) genotypes, the estimated fold change (foldChange), the *P-*value without (pval) and with (padj) FDR correction, gene position on the JGI reference genome (JGI scaffold, start, and end), homologous gene identification number for *Arabidopsis thaliana* (*A. thaliana* ID), *Arabidopsis lyrata* (*A. lyrata* ID), and *Schrenkiella parvula* (*S. parvula* ID), and gene name (Gene name) based on *A. thaliana* ID. Genes identified in both this study and 1) all plants (Shared with Champigny et al. 2013), 2) chamber-grown plants (Shared with Champigny et al. 2013 cabinet-grown plants), and 3) field-grown plants (Shared with Champigny et al. 2013 field-grown plants) are indicated. Genes identified as “True PAVs” were identified as PAVs in both Champigny et al. 2013 and our study with at least 1 read in SH but no read in YK (Yes; SH only) or at least 1 read in YK but no read in SH (Yes; YK only). Genes not identified as PAVs in either study are indicated (No).

**Supplementary Table 2.** Primer sequences used for qRT-PCR. Gene name (Name) and primer sequence (Sequence) for all genes used to test transcriptome expression differences by qRT-PCR.

**Supplementary Table 3.** All metabolites detected in E. salsugineum accessions Shandong (SH), Yukon (YK), and YK x SH F1 hybrids. Data are means (n=4). Empty indicates absence or undetected metabolites. Genotypes are significantly different at P-value ≤ 0.05 (*), 0.01 (**), and 0.001 (***) based on two-tailed t tests in Experiment 1 (E1), Experiment 2 (E2), or nested (N) analysis of the two experiments. In Experiment 2 (E2), S v Y indicates comparison of SH versus YK; F1 v S indicates comparison of YK x SH F1 hybrids versus SH; F1 v Y indicates comparison of YK x F1 hybrids versus YK; F1 v M indicates comparison of YK x SH F1 hybrids versus mid-parent; Nested indicates comparison of SH versus YK for two experiments combined together. Genotypes are significantly different at P-value ≤ 0.05 (*), 0.01 (**), and 0.001 (***) based on two-tailed t test. Different letters within the same line indicate significant difference between genotypes at P-value ≤ 0.05 based on Tukey’s studentized range test. Carbohydrates indicate sugar-related compounds and derivatives in citric acid cycle. Fatty acids indicate fatty acids and their derivatives. Others indicate all other compounds not belong to carbohydrates or fatty acids, or amino acids.

**Supplementary Table 4. Biallelic SNPs between *E. salsugineum* Shandong (SH) and Yukon (YK) accessions.** SNPs between *E. salsugineum* Shandong (SH) and Yukon (YK) identified by our transcriptome sequencing. SNP location is indicated according to the JGI *E. salsugineum* genome by scaffold and position within the scaffold, as well as the gene locus in which the SNP was identified. Reference and alternative alleles are indicated. High quality SNPs, based on our criteria described in the methods section are indicated. SNPs that were identified in this experiment and Champigny et al. 2013 (Shared with Champigny et al. 2013) and/or existing Sanger data (Present in Sanger data) are indicated.

**Supplementary Table 5**. Heterosis effect of metabolites in YK x SH F1 hybrids of E. salsugineum. Metabolites that were different between F1 and mid-parent showed no enrichment in those were different between the two parents based on a χ2 test. Metabolites in F1 hybrids that showed transgressive heterosis were enriched in the metabolites with no variations between the two parents at P-value ≤ 0.05 (*) based on a χ2 test. The proportion of negative heterosis metabolites (23/24) is significantly different from the expected probability of 0.5 based on a Binomial exact test at P-value ≤ 0.0001 (***).

